# Selective enhancement of neural coding in V1 underlies fine discrimination learning in tree shrew

**DOI:** 10.1101/2021.01.10.426145

**Authors:** Joseph W. Schumacher, Matthew McCann, Katherine J. Maximov, David Fitzpatrick

## Abstract

Visual discrimination improves with training, a phenomenon that is thought to reflect plastic changes in the responses of neurons in primary visual cortex (V1). However, the identity of the neurons that undergo change, the nature of the changes, and the consequences of these changes for other visual behaviors remain unclear. Using chronic *in vivo* 2-photon calcium imaging to monitor the responses of neurons in V1 of tree shrews learning a Go/No-Go fine orientation discrimination task, we find increases in neural population measures of discriminability for task-relevant stimuli that correlate with performance and depend on a select subset of neurons with preferred orientations that include the rewarded stimulus and nearby orientations biased away from the non-rewarded stimulus. Learning is accompanied by selective enhancement in the response of these neurons to the rewarded stimulus that further increases their ability to discriminate the task stimuli. These changes persist outside of the trained task and predict observed enhancement and impairment in performance of other discriminations, providing evidence for selective persistent learning-induced plasticity in V1 with significant consequences for perception.

## Introduction

Neurons in primary visual cortex (V1) respond selectively to different stimulus features^1^, constructing neural representations that reliably encode the visual information necessary to support perception and behavior^2, 3, 4^. Visual experience plays a critical role in the early development of these representations^5^, and there is considerable evidence that these representations remain plastic in the mature brain, allowing learning to enhance the discrimination of sensory stimuli necessary to perform novel tasks^6^. This process has been explored through visual perceptual learning paradigms in which behavioral training with visual stimuli results in substantial improvement in discrimination or detection performance^3, 4, 7, 8, 9, 10, 11^. Perceptual learning is thought to result from persistent changes in the activity of a limited population of neurons in the cortical representation whose properties enable the enhanced discrimination capability^11, 12^, but the nature of the changes and the cortical regions associated with such changes are debated^13, 14^. While there is evidence that visual perceptual learning modifies V1 neural response properties such as orientation selectivity ^4, 11^, direction tuning^8^, contrast sensitivity^10^, and contour detection ^15, 16^ many questions remain about how learning-induced changes in neuronal response properties impact the neural representation of relevant stimulus dimensions and how these changes relate to discrimination of trained and non-trained stimuli under active and passive viewing conditions.

To address these issues we employed a 2-photon *in vivo* imaging paradigm that allowed us to longitudinally track the responses of large populations of V1 neurons through an orientation-discrimination learning paradigm in the tree shrew, a close relative of primates with a visual cortex that exhibits a highly-organized functional architecture^17, 18, 19^. With training, animals were able to improve their ability to discriminate the orientation of rewarded and non-rewarded stimuli in a V1-dependent Go/No-Go orientation discrimination task. Enhanced performance was accompanied by changes in the responses of individual V1 neurons that improved the population’s ability to discriminate rewarded and non-rewarded stimuli. These changes were restricted to a select subset of neurons with pre-training tuning properties that were well-suited for identifying the presence of the rewarded stimulus by virtue of preferred orientations that included the rewarded stimulus and orientations biased away from the non-rewarded stimulus. Learning was accompanied by selective enhancement in the response of these neurons to the rewarded stimulus that further increased their ability to discriminate the task-relevant stimuli. Similar, but weaker, learning-related changes were observed under passive conditions, indicating that discrimination learning induces persistent changes with potential to impact perception beyond the trained task. Indeed, in a subsequent modified behavioral task, discrimination of the same rewarded stimulus from neighboring, untrained orientations was enhanced and impaired in ways that could be predicted from the initial training-induced changes in population response. These results suggest that perceptual learning involves persistent changes in the response properties of a task-relevant subset of V1 neurons that modifies the representation of visual stimuli in a way that enhances task performance at the expense of other related discriminations.

## Results

### Tree shrews learn to perform a V1-dependent, fine orientation discrimination task

To assess the role of V1 in learning and performing a perceptual discrimination, we trained tree shrews to perform a simple Go/No Go orientation discrimination task (Fig. 1a). Tree shrews self-initiated individual trials by licking their reward port, and following a variable delay were presented with a 500 ms static oriented grating. If the orientation of the grating matched the assigned rewarded orientation (S+) for the individual shrew, licking during the subsequent 1.5 second response period would result in a liquid reward (hit), while failure to lick (miss) would result in a timeout (see methods and materials). If the orientation of the grating did not match the S+, a lick response (false alarm, FA) resulted in a timeout, while withholding a response was considered a correct rejection (CR) of the non-rewarded orientation (S-). There is considerable variation in the implementation of Go/No Go behavioral tasks for behavioral and neurophysiological studies ^4, 7, 20, 21, 22, 23, 24, 25^, but the typical Go/No Go task has the limitation that non-responses are somewhat ambiguous since they can result from correct task performance or other factors such as attentional lapses or fluctuating motivational factors^7^. Furthermore the timing and ratio of delivery of S+ and S-stimuli can systematically alter task engagement and performance biases^20^. To incentivize sustained task engagement, especially throughout No Go trials, S-CRs were “rewarded” with a subsequent S+ and response period (Fig. 1a). Because these additional presentations had a 100% probability of being the S+, hit responses could be driven by a strategy independent of visual discrimination, and for this reason they were excluded from overall performance calculations. Finally, to provide a disincentive for licking during the stimulus presentation, lick responses that occurred before stimulus offset resulted in aborted trials regardless of trial type (S+ or S-).

**Figure 1:**
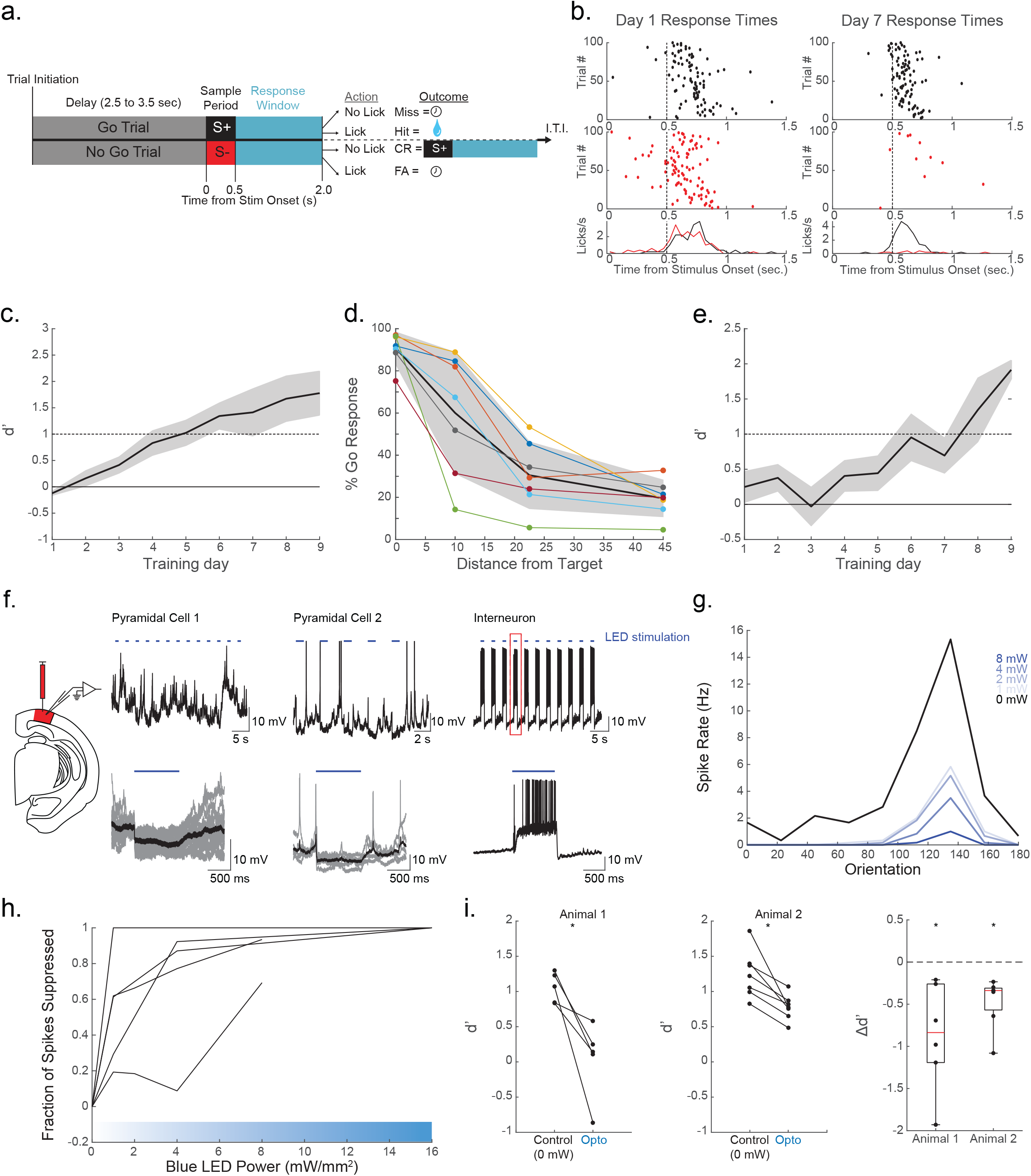
Tree shrews learn to perform a V1-dependent orientation discrimination task. **(a.)** Task structure. Tree shrews self-initiated trials by licking their response port. Following a variable delay, shrews were asked to lick following an S+ orientation while not licking in response to an S-orientation. Hit trials elicited liquid rewards, while correct rejections (CR) resulted in a subsequent S+ presentation. Miss and false alarm (FA) trials resulted in a brief time-out. Inter-trial interval (ITI) was determined by the tree shrew. **(b.)** Representative performance for days 1 (left) and 7 (right) demonstrates coarse discrimination learning. The tree shrew initially licks indiscriminately for both S+ (black) and S-(red) trials. After one week of training the shrew licks predominantly on S+ trials, while rejecting S-trials. Lick time histograms (bottom) show a refinement of response times to immediately follow the stimulus presentation on day 7. **(c.)** Mean (+/- SEM) performance showing shrews on average reached criterion performance within one week of training (n = 11; mean days to criterion = 6.17 +/- 2.92 SD) **(d.)** 7 shrews were trained to perform the task with multiple S-offsets. Performance at 10 degree discriminations was unreliable, while shrews were able to achieve reliable suppression of go responses for S-orientations of 22.5 degrees or higher. **(e.)** Mean (+/- SEM) learning curves show fine discriminations of 22.5 degrees reached criterion levels on average after 7.44 days of training (+/- 3.0) **(f.)** Expression of ChR2 under the mdlx enhancer in V1 enabled blue-light mediated suppression in two example pyramidal neurons (top: raw whole-cell voltage recordings; bottom: individual stimulus locked trials in grey, with average traces in black). Fast-spiking excitation was observed in a putative interneuron (top: raw whole-cell trace; bottom: example stimulus locked trial outlined in red above). **(g.)** Orientation tuning curves recorded at varying levels of blue light power from one example neuron show increasing spike suppression across orientations. **(h.)** 5 neurons showed increasing spike suppression at preferred orientation with as little as 1mW/mm2 **(i.)** 2 headfixed shrews were implanted with bilateral windows over V1 expressing ChR2 via the mdlx enhancer (left). Average discrimination performance in both animals was impaired during trials with optogenetic stimulation in both animals. The left two panels show discrimination performance decrements for individual sessions. The right panel shows Δd’ distributions for each animal.

To become familiar with the testing apparatus and learn the reward contingencies of the task, animals were first trained to perform coarse discriminations (>=45 degrees). Characteristic learning behavior was observed as a refinement of licking response times (Fig 1b). On day 1 (Fig. 1b, left), the animal learns to time licking responses after the 500 ms stimulus presentation, but responds on both S+ (black dots) and S-(red dots) trials, and responses on these trials are distributed over the course of about 1 second (lick timing histograms, Fig. 1b bottom). Following 7 days of training (Fig. 1b, right), the timing of Go responses is more locked to the stimulus onset, and the fraction of responses on No Go trials is substantially reduced, indicating improved discrimination. Tree shrews (n=11) typically reach criterion performance levels (d’>=1) for coarse discrimination in under 7 training sessions (Fig. 1c, left; mean days to criterion = 6.17 +/- 2.92 SD).

To establish the psychophysical limits of behavioral performance under our system, a subset of animals (n=7) were trained to discriminate multiple S-stimuli with offsets from the S+ at increments of 10, 22.5, and 45 degrees. Peak performance at each offset (Fig. 1d) yields individual psychometric curves that demonstrate a group performance breakdown at discriminations of 10 degrees. Animals reliably have high Go rates for S+ stimuli (offset = 0 degrees), and low Go rates for S-offsets from target by 22.5 degrees or higher, but have highly variable performance with 10 degree discriminations, with shrews less able to suppress Go responses. For subsequent experiments, 22.5 degree discriminations were considered “fine” discriminations, because they posed a greater challenge than coarse discriminations of 45 degree discriminations and above, but a majority of animals were able to reach criterion performance at this level of difficulty.

Animals used for the imaging experiments (n = 6) and additional behaviorally trained animals (n = 3) mastered the coarse discrimination task, and were then introduced to an S-stimulus that was offset by 22.5 degrees from the previously trained S+ target stimulus (Fig. 1e; n = 9; days to criterion = 7.44 +/- 3.0 SD). The time to criterion performance did not differ significantly between coarse and fine discrimination periods (p>0.4, two-sample t-test), suggesting that while the coarse task instructed the animal to use the apparatus and learn a set of reward contingencies, there remains a general lack of transference of perceptual skill from coarse to fine discrimination learning.

Before examining the responses of V1 neurons during the discrimination task, we thought it was important to confirm that V1 activity contributes to the performance of this task. To address this, we transiently suppressed V1 activity bilaterally during presentations of the visual stimulus, interleaving trials with or without optogenetic activation of inhibitory interneurons that virally expressed channelrhodopsin^26, 27^ (AAV1.mdlx.ChR2). In separate experiments we validated the suppression of V1 responses with this approach, demonstrating that blue light stimulation induces hyperpolarization of putative pyramidal cells and depolarization accompanied by fast-spiking activity in a putative interneuron (Fig. 1f). We also titrated the intensity of blue light stimulation to arrive at an intensity sufficient to achieve maximal suppression of spike rates (8 mW/mm^2^; Fig. 1g,h). Consistent with the contribution of V1 activity to task performance, in two tree shrews with bilateral mdlx.ChR2 expression, discrimination was significantly impaired on the trials with optogenetic stimulation (Fig. 1i; p<0.01 Wilcoxon signed rank test). We note that the area of inactivation was limited to a small percentage of V1 surface area (estimated from previous reports as approximately 10 to 15%^28, 29, 30, 31^) which likely accounts for the residual performance in these experiments.

### V1 neural populations improve fine discrimination performance with learning

We then asked if the learning of fine orientation discrimination with the Go/No Go task is accompanied by changes in the response properties of V1 neurons that could facilitate task performance, and reasoned that such changes may be reflected in the neural discriminability of task relevant stimuli at the level of the neural population. We chronically imaged neural activity in populations of tree shrew V1 layer 2/3 neurons with two-photon calcium imaging of the genetically encoded calcium indicator GCaMP6s^32^, which we expressed via microinjections of AAV vectors (see methods and materials). This enabled us to measure responses to S+ and S-stimuli during behavioral performance at multiple time points relative to criterion fine discrimination performance (Fig 2a). Alignment of the chronically imaged field of view was facilitated by anatomical landmarks such as neuronal somata, blood vessels, and cortical depth relative to the imaging window coverglass (Fig. 2a, left). Representative example neurons show that raw responsiveness remains relatively stable across learning time points, with notable changes (Fig. 2a,b). Cell 1 is an example of an S+ responsive cell that gains responsiveness over the course of fine discrimination learning, while Cell 2 is an example of an S-responsive cell that has diminished responsiveness across time points (Fig. 2b; for additional examples see Supplementary Figure 1a).

**Figure 2:**
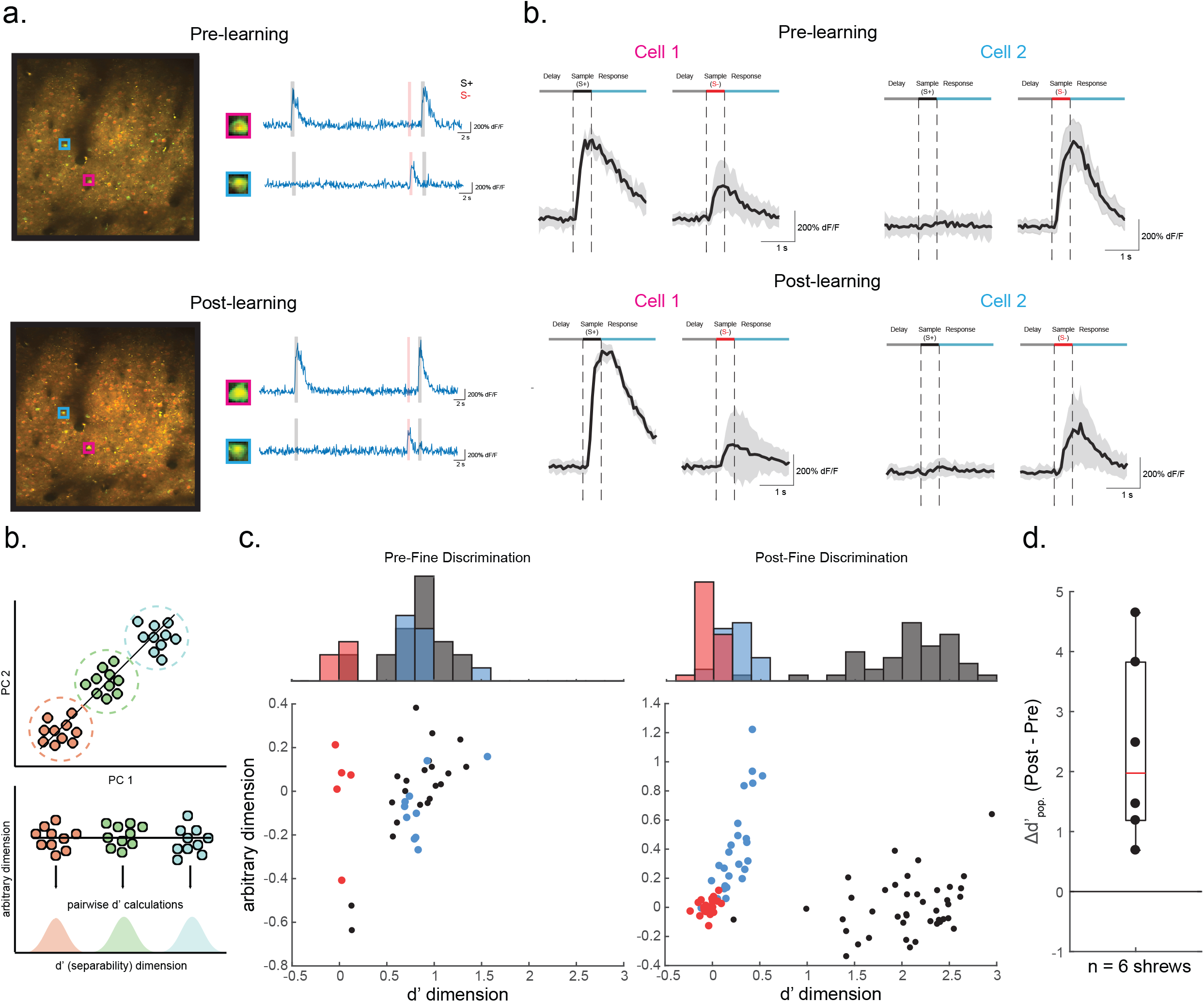
Tracking neural populations over the time reveals learning-related enhancement of neural population discrimination. **(a.)** Neural response properties were measured at pre-(top) and post-learning (bottom) time points during learning. The neurons within the same field of view were identified across sessions. Example neural response traces for tracked cell bodies are shown as raw dF/F traces (right) **(b.)** Mean stimulus locked traces (+/- SEM) for example neurons in **a**.. Cell 1 (magenta) was an S+ selective cell that became more selective over time. Cell 2 (cyan) was an S-selective cell that became less selective over time. **(c.)** Schematic of neural population discrimination metric. Individual population responses are color coded by stimuli and plotted by their first 2 principle components (top). Each data point was then projected onto the line that maximized cluster separability, and pairwise d’ measures were computed for each distribution (bottom) **(d.)** Population responses for an example animal before and after learning in neural discrimination space (bottom) and marginal histogram (top). Pre-learning (left) the S+ (Black) and 22.5 degree S-(blue) are highly overlapping, but are both separable from a 45 degree S-(red). In the post-learning recording (right) the S+ is highly separable from the S-distributions. **(e.)** All six animals display a positive Δd’_pop_. between pre- and post-learning recordings (p<0.05, Wilcoxon signed rank test).

In order to derive a measure of population neural discriminability—i.e., how well the S+ and S-stimuli can be discriminated based on the activity patterns of many V1 neurons, we turned to a dimensionality reduction approach. Each individual trial’s population response (dimensions = n cells) is projected onto a low dimensional space comprised of the first two principle components of the population response (dimensions = 2; see methods and materials). Fig. 2c schematizes this process, where each point represents an individual trial’s population response, and is color coded by the orientation presented on that trial (Fig. 2c, top). To determine the neural population’s capacity to discriminate these clusters of population responses, we calculate the separability of each cluster from one another by projecting each point onto a line that maximizes the distance between each cluster center. Population responses to individual stimuli form distributions of values along this separability dimension (Fig 2c, bottom). These distributions are then the basis on which we quantify neural population discrimination performance (d’_pop._) using a standard separation index (see methods and materials). In the pre-learning phase, this approach demonstrates that the population responses of V1 neurons to the S+ and a coarse (45 degree offset) S-are sufficiently different to allow these stimuli to be discriminated (Fig 2d, left). In contrast, population responses to the S+ and fine (22.5 degree offset) S-for these animals are highly overlapping suggesting that the naïve V1 population responses provide less of a basis for reliable fine discrimination. Following behavioral training (Fig. 2d, right), enhanced behavioral performance of the animals in fine discrimination was accompanied by significant increases in the neural discriminability (p < 0.05 Wilcoxon signed rank test) of these stimuli based on V1 responses within the population of chronically tracked neurons (Fig. 2e).

### Single cell discrimination improvements are orientation specific and linked to reward associations

To better understand changes in the population response of V1 neurons that contribute to enhanced discrimination, we thought it would be important to start by characterizing the patterns of activity that are evoked by the S+ and S-stimuli prior to learning the discrimination. The tree shrew V1 has a well-organized modular architecture for orientation preference^18, 19^, and this was clearly evident in awake head-fixed animals with passive presentation of grating stimuli (Fig. 3a). As expected, presentation of the S+ (Fig. 3b, left) and S-(Fig. 3b, right) stimuli during the behavioral paradigm resulted in robust modular responses that were distinct, but highly overlapping (outlined in 3a), reflecting the orderly progression of orientation preference and the breadth of tuning exhibited by individual layer 2/3 neurons. Well-tuned neurons’ median tuning curve full width at half max = 55 degrees +/- 13.98 SD (see methods and materials), consistent with previous observations in both tree shrew^31^ and primate^33, 34^. This spatial distribution emphasizes that even in the untrained state, individual V1 neurons that are activated by the S+ and S-stimuli differ in the degree to which their activity contributes to discrimination of the two stimuli, depending on the relation of their tuning curve to the S+ and S-orientations ^11, 35^.

**Figure 3.**
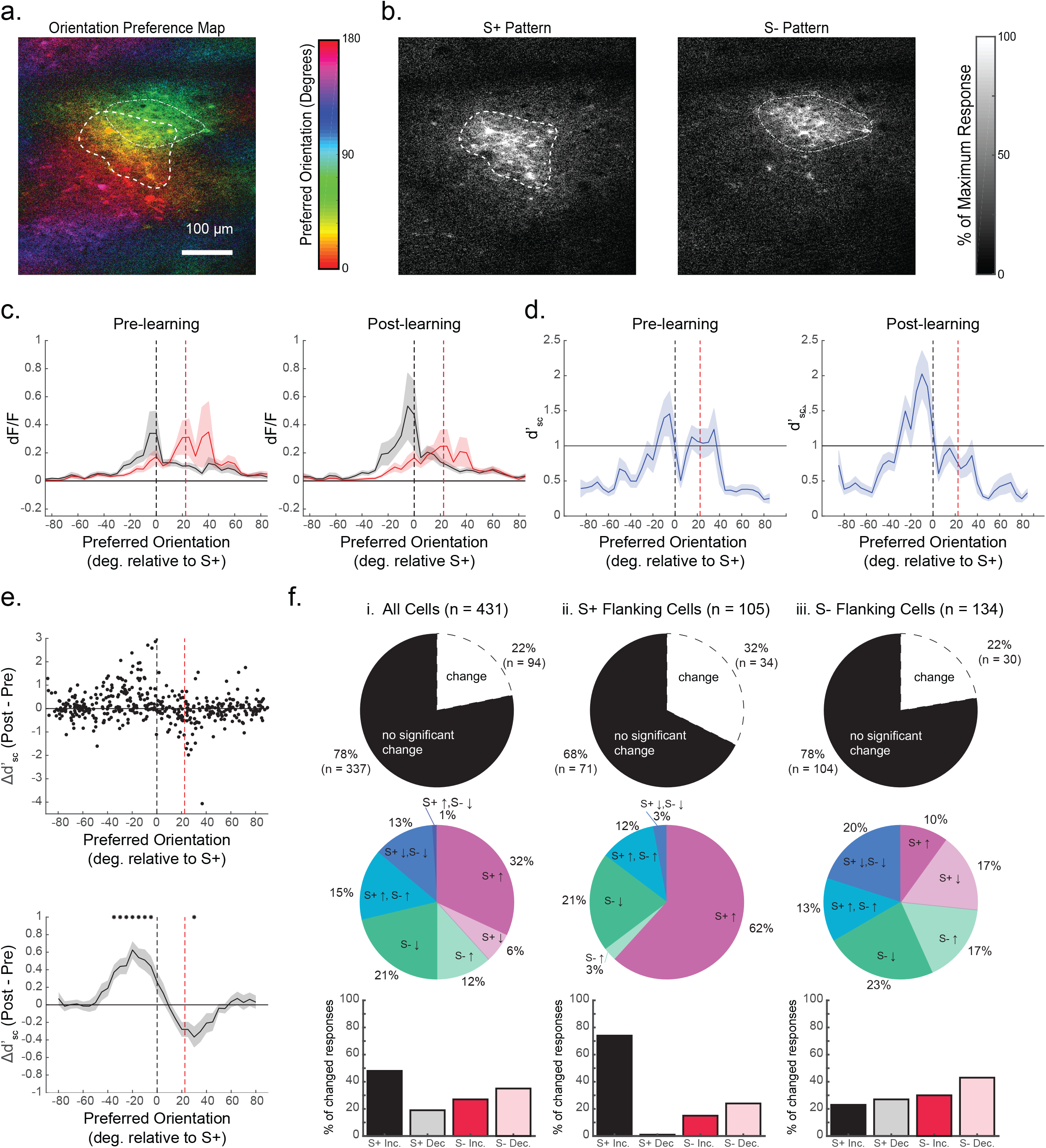
Single cell improvements in neural discrimination are predicted by baseline discrimination capacity and reward associations. **(a.)** Tree shrew V1 contains an orderly map of orientation preference. The orientation preference map is computed from responses to oriented gratings during passive viewing. **(b.)** Average calcium responses to S+ (left) and S-(right) stimuli shown during behavior are distributed over overlapping columns within V1, with many of the same neurons responding for both stimuli. Contours outline cortical regions responding greater than 30% of the maximum response per stimulus, and are overlaid on the orientation map in a. **(c.)** Population response curves from binned single cell responses (dF/F) to S+ (black) and S-(red) stimuli, arranged by preferred orientation for pre-(left) and post-learning (right) conditions. **(d.)** Average binned single cell d’ values arranged by each neuron’s preferred orientation. Pre-learning (left), peak d’ values were observed in neurons with preferred orientations flanking the S+ and S-stimuli, and were lower in between the S+ and S-, indicating that these neurons provide minimal discrimination information for the task. Post-learning (right) a significant bias exists with S+ flanking neurons displaying greater discrimination information than S-flanking neurons. **(e.)** Top: V1 neurons with the greatest learning related improvements in d’_sc_ have preferred orientations neighboring the S+ (black dashed line), but not S-stimulus (red dashed line). Significant increases in Δd’_sc_ were found up to 35 degrees from the S+ stimulus, while one significant decrease in (bottom: averaging delta d’ over a moving window of preferred orientations in 5 degree increments with 20 degree bin size, mean +/- SEM, *p<0.05 Wilcoxon signed rank test with Bonferroni correction). **(f.)** Learning related changes in responses to task-relevant stimuli depended largely on the functional properties of neurons relative to the orientations of the task. Top row: percentages of neurons with a significant change in response to at least one task relevant stimulus out of i. all cells (22%), ii. S+ flanking cells (32%), and iii. S-flanking cells (22%). Middle row: breakdown in specific types of changes observed on a cell by cell basis. Out of all cells and S-flanking cells there were heterogenous changes in the magnitude of responses to both stimuli, while S+ flanking cells largely saw increases in S+ responses. Bottom row: total proportions of significantly changing cells exhibiting increases or decreases in response to the S+ or S-. S+ flanking cells are heavily dominated by increases in response to the S+ (74%), while S-flanking neurons show balanced changes with S-decreases being the prevailing change (43%).

To derive a population response profile that represents the distribution of neural activity evoked by S+ and S-stimuli as a function of a neuron’s orientation preference, passive presentations of the full range of orientations were used to compute the pre-learning preferred orientation for each neuron in our longitudinally tracked sample (see methods and materials). Using this data, cells (n = 431) were binned by preferred orientation (orientation axis normalized across animals: 0 degrees = S+, 22.5 degrees = S-), and a moving average (+/- SEM) across orientation bins (10 degree width, 5 degree increments) was computed for single cell responses to the S+ and S-stimulus (Fig. 3c) In the pre-learning condition (Fig. 3c, left), the peak responses for the S+ and S-stimulus were distributed around neurons with corresponding preferred orientations, but each stimulus evoked responses from neurons with a broad range of preferences, resulting in significant overlap of the two distributions. Following learning (Fig. 3c, right) there was a noticeable change in the distribution of responses to the S+ stimulus, with increases in the responses of neurons with preferred orientations that include the S+ orientation and nearby orientations biased away from the S-orientation. There also appeared to be a modest decrease in the response to the S-stimulus for neurons that prefer the S- and neighboring orientations.

These results suggest that the behavioral increase in discrimination performance is not simply the result of an increase in responses of V1 neurons to the rewarded stimulus, but an increase in response of a select subset of neurons whose tuning is optimal for distinguishing the rewarded stimulus from the distractor. To specifically test the discrimination capability of the population of neurons in our sample, we calculated single cell (S+,S-) discriminability (d’_sc_; separability index, see methods and materials) using single cell *d*F/F signals on a trial-by-trial basis. Then, by arranging baseline (pre-learning) d’_sc_ by preferred orientation and averaging over bins of 10 degrees, we could evaluate the distribution of d’_sc_ values for both pre- and post-learning conditions. As expected, in the pre-learning condition, the distribution of d’_sc_ values exhibits two peaks separated by a trough: the lowest d’_sc_ -values are found for neurons with preferences for orientations in between the two stimuli, while higher d’_sc_ -values are found for neurons that prefer orientations displaced away from this region (Fig. 3d, left)^11, 35^. Interestingly, post-learning, the discrimination profile of the neural population has a strikingly different appearance with an increase in d’_sc_, especially for neurons with preferred orientations neighboring the S+, and biased away from the S-. This is accompanied by a modest reduction in d’_sc_ for neurons with preferred orientations near the S-.

To better understand the learning induced changes in single neuron responses that contribute to the increases in the population d’, we determined the d’_sc_ values for individual neurons pre- and post-learning and computed the difference (Δd’_sc_). Plotting these values according to each cell’s pre-learning preferred orientation (Fig. 3e, top), we confirmed that individual neurons with the greatest improvements in task discrimination performance are those with preferred orientations flanking the S+ orientation and displaced away from the S-orientations (S+ flanking neurons). We quantified this further by computing a moving average of all Δd’_sc_ within 20 degree bins over 5 degree increments. We found significant improvements in Δd’_sc_ for cells with preferred orientations within 35 degrees of the S+ stimulus and shifted away from the S-stimulus, but in no other preferred orientation bins (Fig. 3e, bottom; p<0.05 Wilcoxon signed-rank test with Bonferroni correction). This analysis also revealed a significant reduction in Δd’_sc_ for neurons with preferences near the S-(Fig. 3e, bottom; p<0.05 Wilcoxon signed-rank test with Bonferroni correction).

While the learning induced enhancement in discrimination is clearly biased to the S+ flanking neurons, the single neuron analysis reveals that there is considerable diversity in the behavior of neurons regardless of preferred orientation. To further probe this diversity at the single cell level, we determined the percentage of neurons in our sample that exhibited significant change in response pre- and post-learning, and characterized the nature of this change (methods and materials). In fact, out of all neurons tracked longitudinally (Fig 3f, i.; n = 431), only 22% (n = 94) exhibited a significant change in response magnitude, changes that included increases or decreases in response to the S+ or S- and various combinations. But the distribution of the neurons undergoing change and the types of change they exhibited was distinct for S+ and S-flanking neurons (n = 105 and 134, respectively). Roughly 32% of S+ flanking neurons exhibited significant changes in response, 74% of which exhibited an increase in response to the S+ stimulus (Fig. 3f, ii.). In contrast, only 22% of S-flanking neurons underwent significant changes in response with the greatest fraction (43%) decreasing their response to the S-, and only 23% exhibiting increased responses to the S+ stimulus (Fig. 3f, iii.). These results indicate that while neurons in V1 undergo heterogeneous learning related changes as a group, orientation specific enhancement of neural discrimination is largely driven by increases in S+ responsiveness.

### Learning-enhanced neural discrimination persists outside of task performance and is associated with biased changes in orientation tuning

While these results indicate that training-induced enhancement in neural discriminability can be explained by changes in the response of a select subset of V1 neurons, it leaves open the question of whether this enhancement is context-dependent - only evident during performance of the task - or extends to stimuli presented outside the behavioral paradigm. To probe this issue we examined whether we could detect learning-induced Δd’_sc_ of neuronal responses under passive stimulus presentation conditions (Fig. 4a). As in the behavioral paradigm, we observed (Fig. 4a,b) an overall increase in d’_sc_ that was specific for the S+ flanking neurons (p<0.05, Wilcoxon signed-rank test with Bonferroni correction). These results indicate that fine discrimination learning is accompanied by stimulus-specific increases in d’_sc_ that persist outside of the behavioral context in which they arise. Moreover, as seen with population responses recorded during task performance, in all 6 tree shrews, learning was accompanied by enhancement of passive neural discriminability of the S+ and fine S-(Supplemental Figure 1b,c; p < 0.05 Wilcoxon signed rank test). The consistency of V1 population changes in active and passive contexts suggests that discrimination learning reflects a persistent increase in the discrimination capacity of a subset of V1 neurons whose tuning is well-matched to the task.

**Figure 4.**
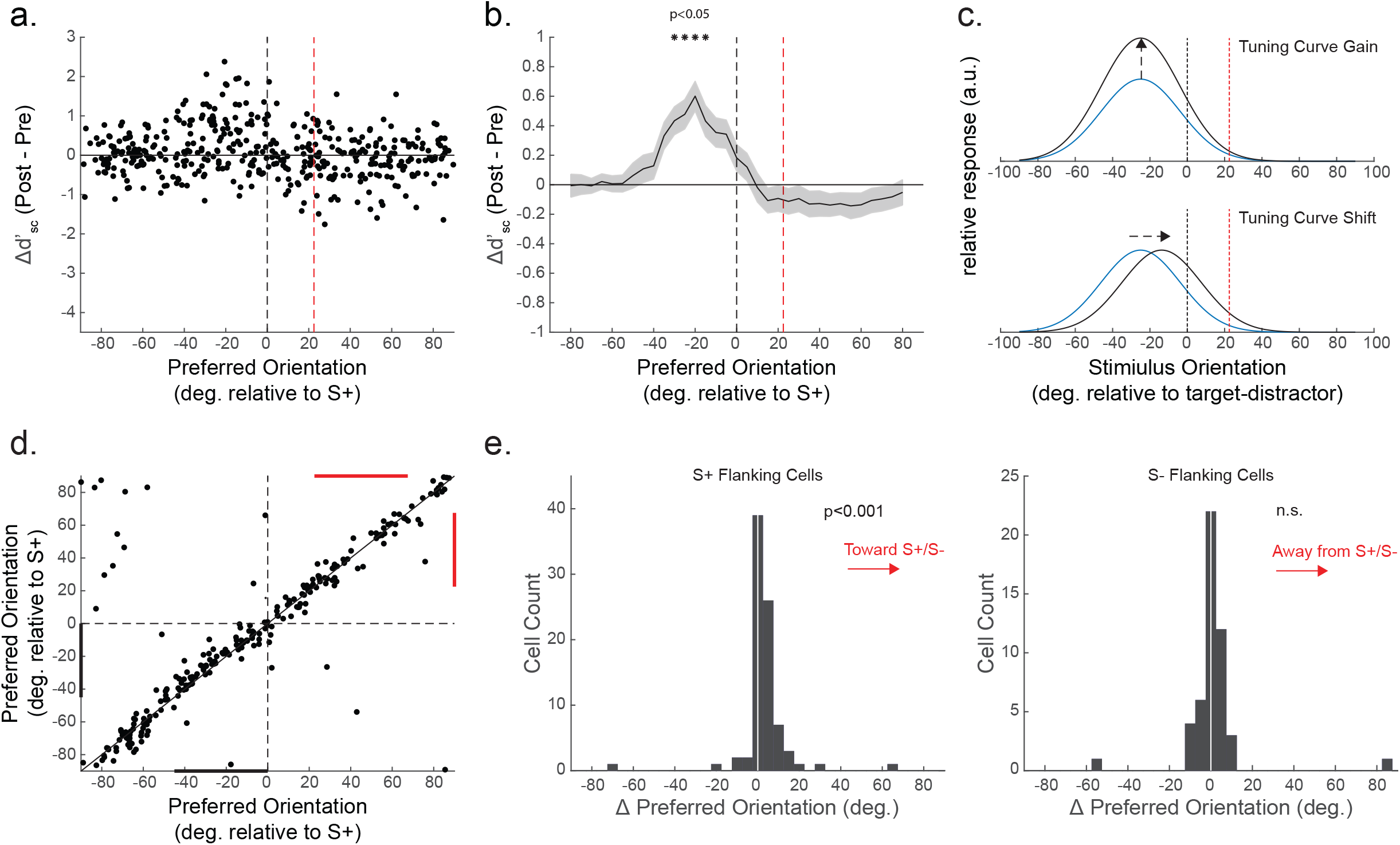
Single cell improvements in neural discrimination persist during passive stimulus presentations and are consistent with biased shifts in preferred orientation. **(a.)** As in Fig. 3e, single V1 cells with the greatest increases in passive d’_sc_ had preferred orientations neighboring the S+, but not S-stimuli. **(b.)** Significant increases in delta d’_sc_ were found between 15 and 30 degrees from the S+ stimulus (averaging delta d’ over a moving window of preferred orientations in 5 degree increments with 20 degree bin size, mean +/- SEM, *p<0.05 Wilcoxon signed rank test with Bonferroni correction). **(c.)** Two types of hypothetical functional changes leading to selectively increased d’_sc_ in V1 neurons. Top: a positive gain in excitation of a single S+ flanking neuron. Bottom: a small shift in preferred orientation toward the S+ orientation. **(d.)** Preferred orientation was remarkably stable in visually responsive neurons before and after fine discrimination learning. Axes are normalized across animals to center the S+ orientation at 0 degrees and the S-orientation at +22.5 degrees. Marginal lines highlight the 45 degrees surrounding the S+ (black) and S-(red) flanks. **(e.)** S+ flanking cells (Left) but not S-flanking cells (Right) exhibit a biased shift in orientation preference (*p<0.001, Wilcoxon signed rank test).

The fact that learning-related effects on neural discrimination persist in the passive condition made it possible for us to further characterize the functional changes in neuronal response underlying the enhanced discrimination of individual S+ flanking neurons. We reasoned that the increased S+ responses in this population of neurons could reflect mechanisms that operate to increase the gain in response, without significant changes in orientation preference (Fig. 4c, Top) or mechanisms that result in a shift in preferred orientation of the cell toward the orientation of the S+ stimulus (Fig. 4c, Bottom). Overall, the pre- and post-learning orientation preferences of the total population of tuned neurons in our sample (tuning curve 1-CV > 0.25, see methods and materials) were highly correlated (Fig. 4d). However, the subpopulation of neurons with preferred orientations in the S+ flank (between −45 and 0) exhibited a distribution of post-learning preferred orientations that were significantly biased toward the S+ relative to the pre-learning state (Fig. 4e, left; positive bias p<0.001, Wilcoxon signed rank test). Significant shifts in orientation tuning (p<0.05, Watson-Williams test for equal means, see methods and materials) were observed in 16.67% of these S+ neurons, while 29.76% saw a significant positive gain in response magnitude at preferred orientation (see methods and materials). The specificity of these changes for the population of S+ flanking neurons is supported by comparable analyses of neurons in the S-flank (between 22.5 and 67.5 degrees). These neurons exhibited no significant directional bias in the distribution of pre- and post-learning changes of orientation preference (p>0.05, Wilcoxon signed rank test; Fig. 4e, right), even with 20.41% of these cells significantly shifting in orientation (p<0.05, Watson-Williams test for equal means). Also, roughly half as many tuned S-flanking neurons saw a significant positive gain at preferred orientation (16.33%) compared to S+ flanking neurons. These results suggest that the increases in S+ responses that drive enhanced discrimination learning reflect small, heterogenous changes in the response properties of V1 neurons tuned near the reward-associated orientation.

### Learning-related changes in population discrimination predict additional behavioral performance impacts

In addition to characterizing the single cell tuning properties that contribute to changes in d’_sc_, the ability to present a broad range of orientations in the passive context allowed us to probe the orientation specificity of the learning-induced changes in neural population discriminability. We computed d’_pop_ between the S+ and orientations within +/- 25 degrees of the S+ in 2.5 degree increments (Fig. 5a). Positive offsets from the S+ indicate orientations in the direction of the trained S-, while negative offsets indicate symmetric, untrained orientations. Consistent with task-specificity for learning-related changes, significant improvements (p<0.05, Wilcoxon signed rank test) in the average (bold trace) population d’ for 6 tree shrews (light traces) reach a peak at 22.5 degrees offset, which is the orientation of the S-(Fig. 5a). Interestingly, enhanced neural population discrimination was observed for untrained orientations that lie between the S+ and S-stimuli (e.g. +12.5 degrees) but not for stimuli with orientations displaced away from the S+ (e.g. −12.5 degrees). Indeed, our analysis reveals weak but significant decreases (p<0.05, Wilcoxon signed rank test) in d’_pop_ for orientations displaced in this direction from the S+. Given that learning-induced improvements in single cell d’ for S+ and fine S-stimuli were localized to neurons with preferred orientations in this range (i.e., S+ flanking neurons) (Fig. 3b), these results raise the possibility that learning may improve the neural discriminability of the trained orientations with consequences for discrimination of nearby untrained orientations.

**Figure 5.**
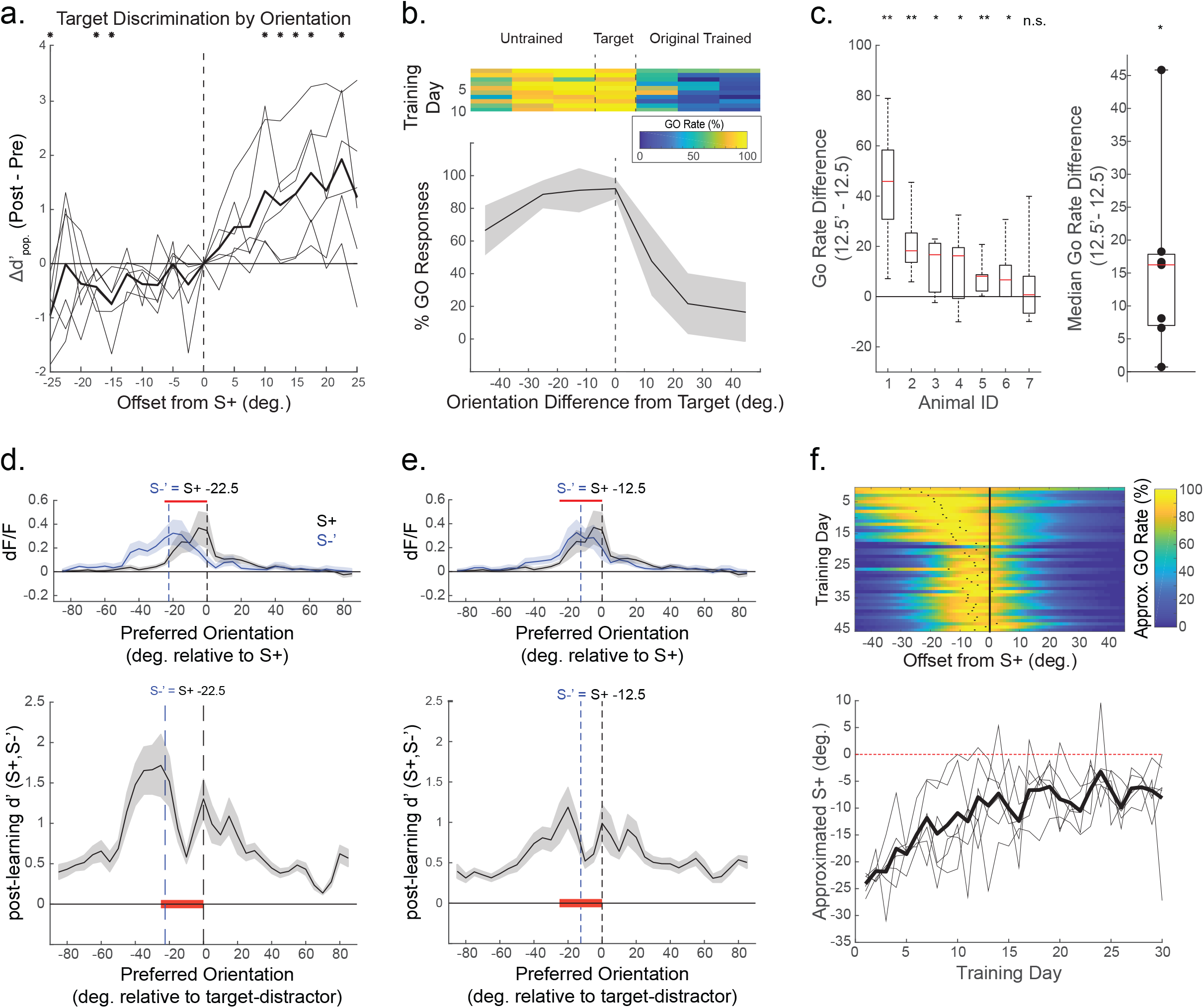
Tree shrew neural discrimination accurately predicts orientation specific benefits and impairments in future behavioral performance. **(a.)** Learning-related changes in d’_pop_ of the S+ (0 degrees) and a wide range of passively viewed orientations further reveals the orientation specificity of the training effect. Across six animal, the peak Δd’_pop_ for population discrimination was seen at the 22.5 degrees (equal to the S-), with gradual increases in d’ observed between the S+ and S-. While no average improvements in d’_pop_ were observed for orientations on the other side of the S+, small but significant decreases in discrimination performance were seen in three of these untrained orientations (*p<0.05 Wilcoxon signed rank test). **(b.)** Tree shrews that initially learned the original asymmetric discrimination were introduced to a generalized, symmetric discrimination task in which multiple S-stimuli were introduced on either side of the S+ orientation. 10 days of performance after tree shrews were introduced to a novel task shows that two example shrews were biased to go for novel orientations, but maintained accurate performance for the original discriminations (>= +22.5 degrees). **(c.)** 6 out of 7 tree shrews showed significant transference of skill in discriminating the S+ from novel +12.5 orientation compared to the novel −12.5 S-orientation (Left: **p<0.01, *p<0.05, Wilcoxon signed rank test). Across animals (right), median go rates were higher for the −12.5 degree compared to the +12.5 degree S-stimulus (p<0.05, Wilcoxon signed rank test). **(Top d**.,**e.)** Passively recorded population response curves (as in Fig. 3c) for the S+ (black) and a hypothetical novel S-(**d**.: −22.5; **e**.: −12.5; blue) after shrews were trained on the original fine discrimination. Regions of the population that are enhanced through reward association (red) overlap with the regions of the population that respond robustly to both the S+ and novel S-**(Bottom d**.,**e.)** Average post learning d’ for the novel S-’ and original S+ arranged by preferred orientation. Regions with poor discrimination are overlapping with the subpopulation of neurons that are enhanced through reward association for the original task, predicting that the learning of this novel discrimination should be impaired following learning of the original fine discrimination task. **(f.)** Gaussian curves were fit to each day’s psychometric function (see **5b**) and tracked for at least 30 days. The peak of the Gaussian is taken as an approximation for the orientation at which the shrew exhibits it’s peak go rate, and thus represents the orientation at which the shrews exhibit minimal discrimination from the S+. Over 30 days of behavioral training, shrews maintain a persistent impairment in discriminating stimuli on the novel side of the S+, and on average continue to associate stimuli within approximately 10 degrees of the S+ with a go response.

If the passive increases in neural discrimination reflect changes in V1 activity that support enhanced behavioral performance, this predicts an asymmetric transfer of discrimination enhancement: tree shrews that have learned 22.5 degree discriminations would show an enhanced ability to discriminate the S+ stimulus from untrained +12.5 degree stimuli compared to −12.5 degree stimuli. To test this prediction, we introducing a novel task to shrews that had been trained on the original S+/S-discrimination. Following the completion of combined imaging and behavioral experiments, tree shrews continued to train in their home cages for at least 10 days in the original task. We then modified the task to include S-stimuli at symmetric offsets from the S+ stimulus at +/- 12.5, 22.5 and 45 degrees. Figure 5b shows the average performance of one shrew on the symmetric Go/No Go task over an initial 10 day period. Consistent with the prediction, there was a conspicuous asymmetry in discrimination performance, with a much lower percentage of Go responses (i.e. correct rejections) for S-stimuli offset by +12.5 degrees from the S+ stimulus (in the direction of the original S-stimulus) than were found for stimuli offset by − 12.5 degrees from the S+ stimulus (in the direction away from the original S-). Significant differences in go rates at +12.5 and −12.5 degrees were observed in 6 out of 7 shrews (Fig 5c, left), and as a group median go rate differences were significantly different from zero (Fig 5c, right; p<0.05 Wilcoxon signed rank test).

While this asymmetry is consistent with a neural enhancement in discrimination performance for untrained stimuli offset by +12.5 degrees from the S+, we recognized that additional factors may contribute to the high percentage of incorrect responses to untrained stimuli offset by −12.5 from the S+. Indeed, our results show that the neurons best able to discriminate the S+ from the original S-stimuli—the S+ flanking neurons—are especially poor at discriminating the S+ stimulus from nearby stimuli offset in this negative direction (Fig 5d,e). Thus, if the animals have learned to rely on the activity of S+ flanking neurons to signal the presence of the S+ stimulus (i.e., the reward), and these neurons respond in a similar fashion to the S+ stimulus and these neighboring stimuli, poor discrimination performance for these stimuli would be expected. Indeed, not only did animals exhibit an initial asymmetry in discrimination performance, the high Go-rate for negative offset stimuli continued even after 30 days of training, well beyond the normal range of time to reach criterion performance for the original S+/S-stimuli (approximately 1 week; Fig 5f). Each shrew’s approximated S+ stimulus was estimated by taking the peak of a Gaussian fit of each day’s psychometric curve (Fig. 5f, top). While there was progressive improvement over time in the Go rate (false alarm rate) for negative offset stimuli at 45 and 22.5 degrees from the S+ stimulus, these improvements reached a plateau for the 12.5 degree offset stimulus (Fig. 5f, bottom) − the stimulus predicted to be the least discriminable from the S+ stimulus since it drives a neural population response that has the highest degree of overlap with the population that signifies the presence of the S+.

Together, these results suggest that while Go/No Go training is accompanied by changes in V1 responses that optimize discrimination of the rewarded and distractor stimuli, these changes could be responsible for an orientation specific impairment in discrimination learning of nearby orientations that persists over several weeks. Thus persistent learning-induced changes in the responses of V1 neurons that are optimized for performance of one behavioral task, may be counter-productive for learning to discriminate other similar visual stimuli.

## Discussion

Here we demonstrated that learning a fine orientation discrimination task is accompanied by persistent changes in the responses of neurons in layer 2/3 of tree shrew V1 that enhance the neural discriminability of task-relevant orientations. This enhancement arises in a select subset of neurons whose preferred orientation is offset from the target stimulus which, prior to training, endows them with a relatively heightened capacity to discriminate the rewarded and non-rewarded stimuli. With learning, the responses of these neurons to the rewarded stimulus increase, further enhancing the neural discriminability of rewarded and non-rewarded stimuli and doing so in a fashion that selectively augments the population response to the rewarded stimulus. The persistence of these changes outside of the task in which it originated, and the enhancement and impairment of learning in additional discrimination tasks that are predicted by these changes, provide converging lines of evidence that plastic changes in the neural representation of stimulus features by V1 neurons contribute to fine discrimination learning.

The demonstration that neurons in tree shrew V1 can exhibit changes in response to task-relevant stimuli is consistent with previous studies that have shown changes in the response properties of V1 neurons that correlate with performance ^3,4,6, 9, 36, 37, 38^. Many of these studies in the rodent have emphasized that it is the increased response of V1 neurons to the rewarded stimulus (and in some cases the non-rewarded stimulus) that correlates with improved task performance. We also find that an increased response to the rewarded stimulus is the prevailing feature of the learning induced change (found in 74% of altered S+ flanking neurons), followed by a reduction in response to the distractor stimulus (found in 43% of altered S-flanking neurons). But a critical aspect of the learning-induced change that is missed in just considering the changes in response to S+ and S-is how these changes impact the population code for discriminating the two stimuli. Our results show that learning is accompanied by a dramatic increase in the ability of V1 neurons to discriminate target and distractor stimuli, and that this is due to a bias in the enhancement of S+ responses to a subset of neurons, S+ flanking neurons, whose tuning at the onset of training is well suited for discriminating the task stimuli. The idea that training selectively enhances the responses of V1 neurons whose tuning profiles make them optimal for discrimination of task stimuli has been suggested before in an orientation discrimination study in primates^11^, but subsequent work did not find any changes in the orientation tuning of V1 or V2 cells in a similar paradigm^13^. However, both research groups reported learning-related orientation tuning changes in V4^12, 14^. By tracking the responses of individual neurons over the course of learning, our data supports that idea that a V1 with highly organized functional architecture undergoes orientation-specific changes during perceptual learning that are constrained by the information coding properties of neurons relative to the task. However, it remains highly likely that V1 is one of many cortical regions that are relevant for perceptual learning.

At the same time, preexisting discrimination capability for task-relevant stimuli does not fully capture the properties of the V1 neurons that undergo learning-induced changes; if this were the only factor, we would expect to see comparable enhancement of the responses of neurons flanking the S-stimulus to the presentation of the S-stimulus. In fact, while S- and S+ flanking neurons exhibit comparable discrimination of S+ and S-stimuli at the start of training, post-learning, the responses of S-flanking neurons actually exhibit reduced discrimination of the task stimuli, in part due to a reduction in response to the S-stimulus. Thus, learning in this task appears to selectively enhance the response of neurons with tuning properties that can optimally discriminate the task stimuli and that respond preferentially to the rewarded stimulus. In short, learning amplifies the responses of neurons that reliably predict the presence of the rewarded stimulus.

Of course, it remains to be seen how much the specifics of the task impact the types of learning induced changes that are present in V1. For example, the task employed here requires discrimination of orientations that activate overlapping distributions of neurons and the neurons that are optimal for predicting the presence of the rewarding stimulus are those that have orientation preferences in the S+ flanks. For performance of Go/No Go tasks that do not require fine discrimination (e.g. 90 degree offset) reliable prediction of the rewarding stimulus could be achieved by enhancement of a population centered on the neurons that prefer the rewarded stimulus, consistent with results in rodents^4, 9 36^. Also, the Go/NoGo task employed here is structured so that only one of the stimuli is associated with reward, and enhancement of a single population of neurons achieves both reliable discrimination and identification of the rewarded stimulus. If instead the task was structured so that both stimuli were associated with reward, and discrimination was necessary to determine the appropriate action required to achieve reward (e.g.,2-alternative forced choice, A/B, or same/different tasks) the changes might involve enhancement of two populations of neurons with preferred orientations arranged symmetrically around both stimulus flanks. It remains to be seen just how flexibly the circuits in V1 can be altered to enable different types of discriminations under different task conditions.

Our results demonstrate that the selective enhancement of the responses of V1 neurons is not confined to the behavioral paradigm, but is maintained under passive viewing conditions. Such persistence would be expected if the changes in V1 contribute to the improvement in task performance that persists across days and weeks. This also indicates that the changes we see reflect fundamental alterations in the representation of visual stimuli by V1 neurons, rather than context-dependent neuromodulatory effects, which are known to regulate neural responses in V1^39, 40, 41^. Of course, this does not dismiss the possibility that context-dependent modulatory effects contribute to the effectiveness of task performance, and indeed, the enhancement effects in V1 under the passive presentation conditions are weaker than those observed during active participation in the task. This suggests that the changes we see in V1 responses coordinate with behaviorally driven modulatory effects to enable highly reliable behavioral performance.

Further evidence for persistence of the changes in V1 responses beyond the behavioral paradigm in which they arose emerged from additional experiments where animals were required to discriminate the S+ stimulus from nearby orientations with either a positive offset (towards the S-) or a negative offset (away from the S-). Although perceptual learning effects are generally regarded as being highly selective for the learned stimuli^4, 8, 9, 42^, the persistent learning induced changes in neural population response that we found predicted impacts on these novel discriminations—both enhancement and impairment—that were evident in the animals’ behavioral performance. The enhancement in the acquisition of discriminations for positive offset orientations is consistent with the persistent enhancement in the discrimination capability of S+ flanking neurons and the fact that analysis shows that this extends to the novel negative offset stimulus orientation. Thus, the activity of the S+ flanking neurons continues to be highly predictive of reward, responding strongly to the S+ stimulus and weakly to both the S- and the novel nearby orientation.

The impairment in the discrimination of the S+ stimulus from nearby stimuli with negative offsets is also predictable from the persistent enhancement in the response of S+ flanking neurons to the S+ stimulus since the enhancement reduces the difference in the pattern of population activity evoked by the S+ stimulus and these nearby orientations. This result is reminiscent of the classical psychophysical phenomenon known as the peak shift effect, in which the learning of S+/S-discriminations of a broad range of multisensory features leads to a heightened response rate for untrained stimuli neighboring the S+^43, 44^. In psychological terms, this effect is considered a form of generalization learning, in which trained stimulus attributes are generalized across a range of untrained stimulus features. Our results suggest that what has been described as a generalization of the S+ stimulus to neighboring orientations is, in fact, a byproduct of a learned, neural specialization for discrimination of the original S+ and S-, which results in impairment of discriminations between the S+ and nearby orientations with positive offset.

But what accounts for the persistent impairment in the animals’ ability to learn to discriminate the S+ stimulus from these nearby orientations in spite of extensive training? Given that changes in neuronal response that were associated with learning of the initial task reduced the neural discrimination of the S+ and these nearby orientations, additional changes in neuronal response are likely to be necessary to achieve reliable behavioral discrimination of these stimuli. These changes would include a reduction in the response of the S+ flanking neurons to the S+ stimulus, and an increase in response to the S+ stimulus by neurons with preferences biased towards the original S-stimulus. Both of these changes would have the effect of reducing the neural discriminability of the S+ stimulus from the S- and negative offset stimuli, and reduce the overall reliability of the V1 signal in predicting the presence of the rewarded stimulus in the task. This is because the persistent changes from learning the original task should enable the activity of the S+ flanking neurons to continue to be a good over-all predictor of reward in the second task, discriminating the S+ stimulus from the S- and negative offset stimuli, even without accurate discrimination of the positive offset stimulus. These considerations emphasize that the alterations in the responses of V1 neurons to achieve fine discrimination learning demonstrated here have limits that are ultimately defined by the orientation tuning selectivity of individual neurons and by the training context in which the stimuli are presented. Thus, circuit level changes that enable enhanced population coding necessary for one discrimination can be detrimental to others, the neural population equivalent of a zero-sum game.

While we have focused on the net enhancement of the V1 responses to the S+ stimulus as the principal learning induced change in V1 response, our chronic analysis of single cell behavior through the course of training reveals a diverse set of changes in responses to the S+ and S-stimulus with overlapping but distinct profiles for changes in responses of neurons that prefer orientations in the S+ and S-flanking regions. Understanding how this diversity of changes is achieved at a circuit-level will be an important challenge. Experience-dependent changes in V1 may depend on modulated activity from a wide range of top-down and/or reciprocally connected cortical areas^45, 46, 47^, that are known to undergo learning-related changes, and reward signaling and reinforcement is likely to be further mediated by neuromodulatory inputs which are known to mediate attention and learning in V1^39, 48, 49^. Perhaps importantly, the majority of the neurons in our sample exhibited no significant change in response during the course of learning, suggesting that the V1 plasticity mechanisms responsible for learning engage cortical networks with a degree of specificity that extends beyond the orientation preferences relevant for the task discrimination. Further studies exploring the identity of the neurons that undergo learning induced change, their patterns of connections, and the synaptic mechanisms responsible for the changes in their responses to visual stimuli are necessary to better understand how these specific changes in V1 responses contribute to training enhanced perceptual discrimination.

## Methods and Materials

All experimental procedures were performed in accordance with NIH guidelines and were approved by the Max Planck Florida Institute for Neuroscience Institutional Animal Care and Use Committee. Tree shrew (*Tupaia belangeri*, n = 16, approximately 6 − 36 months of age, male and female) numbers were minimized to conform to ethical guidelines. Of the animals included in this study, 6 were included in combined imaging and behavioral experiments, two were included in electrophysiological verification of optogenetics, two were included in the optogenetic perturbation of behavior, and 6 provided additional behavioral data.

### Surgery and Viral Expression

Tree shrews were first anesthetized with Midazolam (5 mg/kg, IM), Ketamine (75 mg/kg, IM). Atropine (0.5 mg/kg, SC) was administered to reduce secretions, Dexamethasone (1 mg/kg, IM) was given to reduce inflammation during surgery, and Buprenorphine SR provided a long lasting analgesic for post-operative recovery. The animal’s head was shaved, and any remaining hair was removed with Nair. The surgical site was injected with a mixure of bupivacaine and lidocaine (0.3 – 0.5 ml, SC). A mixture of oxygen, nitrous oxide (O2/N20 1:0 to 1:2) and gas anesthesia (isoflurane 0.5 to 2%) were initially delivered through a mask and later switch to an intubation tube. Venous cannulation (tail or hind limb) and tracheal intubation were established after the animal no longer responded to a toe-pinch. Internal temperature was maintained by thermostatically controlling a heating pad. Expired CO2 and heart rate were monitored for any signs of stress. The respiration rate (100 to 120 strokes per minute) was regulated through a ventilator. The animal was placed in a stereotaxic device (Kopf, Model 900 Small Animal Stereotaxic Instrument). A small incision was made in the scalp, and skin and muscle were retracted. After cleaning the underlying skull, the metal headpost and cranial imaging chamber (centered over V1) were affixed to the skull metabond (C & B), and dental acrylic (Ortho-Jet, Lang). The skin and muscle were then unretracted, cut to overlap with the edge of the dental acrylic, and secured in place with vetbond. A circular craniotomy (approx. 6mm diameter) was performed in the center of the cranial imaging chamber with a small drill to expose the dura. In some cases the headpost was implanted separately from the imaging chamber, but the imaging chamber was always implanted along with viral injections.

Visual cortex was injected with a virus expressing GCaMP6s (*AAV9*.*Syn*.*GCaMP6s*.*WPRE*.*SV40*, Penn Vector Core; *AAV1*.*hSyn1*.*mRuby2*.*GSG*.*P2A*.*GCaMP6s*.*WPRE*.*SV40*, Addgene) at 3 to 5 sites (1-2 μl; 1E13 GC/ml) through a beveled glass micropipette (15 to 25 μm tip diameter, Drummond Scientific) using a pressure injector (Drummond Nanoject II), at 200 and 400 μm from the cortical surface. After a brief waiting period, a durotomy was performed within the craniotomy, and a double-layered cover slip composed of a small round glass coverslip (3 – 5mm diameter, 0.7mm thickness, Warner Instruments) glued to a larger coverslip (8mm diameter, 0.17 mm thickness, Electron Microscopy Sciences) with an optical adhesive (Norland Optical Adhesive 71) was placed into the chamber with the thick coverglass gently resting on the brain). The top layer of coverglass was held in place with a snap ring (5/16” internal retaining ring, McMaster-Carr) and sealed with a layer of Vetbond. After sealing the imaging chamber, Neosporin was applied to the wound margin and animals recovered from anesthesia on a heating blanket. Post-operative care included antibiotics (Baytril, 5 mg/kg), and following the timecourse of Buprenorphine SR, anti-inflamatories (Metacam, 0.5mg/kg).

### Two-photon calcium imaging and data processing

Imaging experiments were performaed using a Bergamo II Series Microscope (Thorlabs) using 920 nm excitation provided by a Mai Tai DeepSee laser (Spectra-Physics) running Scanimage 2015 or 2018 (Vidrio Technologies) and an FPGA module (PXIe-6341, FlexRIO, National Instruments). Average excitation power at the objective (16x, CF175, Nikon Insttuments) ranged from 40 to 100 mW. Images were acquired at 15 Hz (512×512 pixel field of view ranging from 1.19 to 1.85 µ/pixel) Two-photon frame triggers from Scanimage and events denoting visual stimuli and phases of behavioral trials were recorded using Spike2 (CED, Cambridge, UK).

In 6 tree shrews, imaging was carried out across training sessions. Prior to data acquisition, the imaging site was located by matching the FOV to known anatomical features from prior recordings, such as blood vessel patterns and somata locations.

### Visual stimulation

Visual stimuli were displayed on a LED monitor with a resolution of 1920 × 1080 pixels and refresh rate of 120 Hz, which was centered in front of the animal at a distance of 25 cm from the eyes. Stimuli were generated using Psychopy2 written in Python.

### Behavioral training

Behaviorally-trained animals had a normal diet consisting of food pellets and supplemental fruits, vegetables, and mealworms. Tree shrews were shaped and trained to perform a coarse visual discrimination task (>=45 degrees) over the course of one to two weeks before learning to perform a fine discrimination task (22.5 degrees). Initial shaping and familiarity with the apparatus was achieved in freely moving animals in a behavioral annex that was either attached to their homecage or attached to the imaging table. The animals were gradually acclimated to handling, weighing, transportation between the animal facility and imaging, and learning to lick a response port for liquid reward over the course of weeks. Liquid rewards were RO water or a 1:1 dilution of apple juice in RO water depending on animal preference. Reward delivery was controlled by a syringe pump with custom electronics (custom parts or BS-8000, Braintree Scientific) through a gavage needle that acted as a capacitive sensor for lick detection through a custom Arduino Uno interface. Animals that showed a willingness to acquire liquid rewards in the absence of a task in the imaging room were acclimated to head fixation and tube restraint over several days in increasingly long durations starting with under a minute and lasting up to 30 minutes. Animals that showed a willingness to acquire liquid rewards while head-fixed were candidates for visual discrimination learning and imaging. Some animals that were not used in head-fixed experiments were able to perform in the freely-behaving setup, and are included in the characterization of behavioral performance (Fig. 1).

During behavioral training, a trial started when the tree shrew licked the reward port. Following a variable delay period (2.5 to 3.5 seconds), the S+ or S-was presented for 0.5 seconds (90-100% contrast, static square wave grating). To match V1 sensitivity, the spatial frequency of the gratings was 0.25 cycles per degree (Lee, et al., 2016). To ensure that orientation was the only feature distinguishing the S+ and S-, grating phase was randomized. Following presentations of the S+, licking in the 1 second response period elicited a liquid reward (hit trial), and failing to lick resulted in a time out (miss trial). Following presentations of the S-, licking in the response period elicited a time out (false alarm, FA), while withholding a response (correct rejection, CR) queued up a subsequent S+ presentation and response period. These guaranteed S+ stimuli were not factored into the quantification of the animals behavioral performance, but rather incentivized CRs. Behavioral performance was measured using behavioral d-prime, d’_b_ = Φ^-1^(Hit Rate) − Φ^-1^(FA Rate), where Φ^-1^ is the normal inverse cumulative distribution function (Poort et al., 2015). Early on in training, the timing of responses was shaped by delivering a small bolus of liquid reward on a subset of trials (hints). As response timing improved, the probability of hinted trials was decreased gradually to zero across sessions.

### Electrophysiology

To evaluate the efficacy of optogenetic interneuron stimulation, whole-cell patch clamp and juxtasomal recordings were performed in two anesthetized tree shrews expressing ChR2 (under control of the mDlx enhancer) by inserting a pipette through an agarose-filled craniotomy covered with a coverglass with a small hole drilled for pipette access. These procedures have been described elsewhere (Wilson et al., 2018) but briefly, a silver-silver chloride reference wire was inserted below the muscle. Recordings were made in current clamp mode using custom Labview software. Pipettes of 5 – 9 MOhm resistence were pulled using borosilicate glass (King Precision Glass) and filed with an intracellular solution containing (in mM) 135 K gluconate, 4 KCl, 10 HEPES, 10 Na_2_-phosphocreatine, 4 Mg-ATP, 0.3 Na_3_GTP, pH 7.2, 295 mOsm. Neurons were recorded from layer 2/3 (100 to 300 um below the cortical surface). Using a Multiclamp 700b (Molecular Devices). Series resistance and pipette capacitance were corrected online, and analog signals were digitized using Spike2 (CED). For optogenetic inactivation, a fibre (1mm, NA 0.63) coupled to a 455 nm LED light source (Prizmatix) was lowered to 3mm above the cover glass. Light power at the cortical surface varied from 1 to 16 mW/mm^2^. Optogenetic stimulation coinciding with visual stimulation started with a brief ramp (100ms) before visual stimulation.

Recordings took place after two weeks of viral expression. Viral injections for acute procedures in tree shrews are similar to those for chronic procedures described above and have been described previously (Lee et al., 2016), but don’t involve implantation of a head-post or chamber, and require only small burr holes rather than a full craniotomy. For electrophysiological recordings, tree shrews were first anesthetized with Midazolam (100 mg/kg, IM) and Ketamine (100 mg/kg, IM) and atropine (0.5 mg/kg, SC) was administered to reduce secretions. A peripheral venous cannula was inserted in the hind limb to allow fluids delivery (10% dextrose in LRS) during surgery. The incision site was treated with a mixture of lidocaine and bupivacaine (0.3 – 0.5 ml, SC) and ear bars were coated with lidocaine ointment (5%). Gas anesthesia was administered as described above in the surgical procedures, and maintained throughout the duration of the craniotomy procedure and experiment. The recording chamber was identical to our typical imaging chamber described above, with the addition of metal extension for head fixation.

### Optogenetic inhibition of bilateral V1 activity

One male and one female adult tree shrew were used for combined visual discrimination behavior and optogenetic stimulation. The shrews were first trained to perform the visual discrimination task to proficiency (d’ > 1). Headpost, window implantation and viral expression procedures were the same as described for GCaMP6s experiments with slight modifications to allow for bilateral windows. Rather than implanting a metal chamber 5mm diameter circular craniotomies were made over bilateral V1 using a 5mm disposable biopsy punch (Integra Miltex). A window comprised of one 4.5mm coverglass (0.7mm thick) was affixed to a wider 4.5mm coverglass (0.17mm thick) with optical adhesive, and following a full durotomy the thick coverglass was placed in the craniotomy on the surface of the brain. The coverglass was sealed in place with vetbond and dental acrylic.

After recovery from surgery, tree shrews resumed behavioral practice with interleaved trials of light stimulation that ramped up and down 100 ms before and after stimulus onset.. Optical fibers were inserted into 3d printed black plastic cylinders that were fit to the surface of the cranial windows to provide sealed light stimulation, support the optical fiber at a fixed distance from the surface of the glass, and ensure uniform illumination. Light intensity was calibrated using both thermal and photodiode light power sensors (Thorlabs), and ranged from 0 to 8 mW/mm^2^.

### Data analysis

Imaging data were first corrected for motion and drift using customized image registration software written in Matlab (Mathworks). Cellular regions of interest (ROIs) corresponding to visually identified neurons were assigned using ImageJ by either inspecting frames of an imaging stack, or the average intensity or standard deviation z-projections of the imaging stack. The fluorescence time series of each cell was measured by averaging all pixels within the ROI over time. Evoked fluorescence signals were calculated as *d*F/F=(F−F_0_)/F_0_, where F_0_ is the baseline fluorescence signal averaged over 0.5 seconds before stimulus onset, and F is the average fluorescence signal during the stimulus presentation.

Orientation tuning curves were measured by calculating the mean *d*F/F for each orientation. Preferred orientation was calculated by computing a vector sum of the tuning curve and measuring angle of the resultant vector in polar coordinates (CircStat Toolbox^50^). Tuning curve bandwidth was calculated by first least-squares fitting a double Gaussian to the tuning curve, and taken as the full width at half max of the Gaussian function to the nearest degree. Single neurons were defined as orientation tuned based values of 1 – circular variance (CV) > 0.25 which was defined as

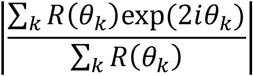

Where *θ*_*k*_ is the orientation of a visual stimulus and R(*θ*_k_) is the response to that stimulus. Statistical significance of shifts in orientation preference were measured with a Watson-Williams two sample test for equal means in circular data (CircStat Toolbox^50^). Statisitcal significance of changes in S+, S-, or preferred orientation response magnitude were calculated using Wilcoxon rank sum tests between pre- and post-response distributions.

We quantified the neural discriminability of pairs of behaviorally relevant stimuli at the neural population level using a population d’ neurometric (d’_pop._). This metric relies on PCA to reduce the complexity of the simultaneous responses of many cells to a lower dimensional space in which separability analyses can be performed. First, the population response on each trial is organized an nx1 dimensional vector of ΔF/F, where n is the number of cells in the neural population. Population responses across trials and stimuli are then arranged into an mxn matrix, where, m is the number of trials across stimuli. For the analysis of d’_pop._ of the active (Fig. 2c,d) and passive responses (Supplemental Fig. 1b,c) to the PC space is computed using responses to the orientations that bound the behaviorally relevant orientations (S+ and coarse S-orientations). Principle component coefficients and scores are computed for the m-by-n population response matrix, and individual trials are projected on to the first two principle components. Individual trial points are then projected onto the 1d space defined by the line that connects the mean responses of individual stimuli. Neural discriminability between pairs of stimuli is then quantified as the d’ of distributions of single trial responses in this 1d space,

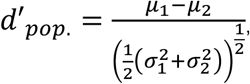

where µ_1_ and µ_2_ were the mean position in the 1d space, and σ_1 and_ σ_2_ were the standard deviations of those posltions. Using passively recorded responses, we also analyzed d’_pop._between the S+ and other orientations spanning +/- 25 degrees around the S+ (Fig. 5a). Here, the PC space was fit using responses to the S+ orientation and either positive or negative 25 degrees from the S+, depending on the offset of the paired orientation from the S+.

We quantified the neural discriminability of pairs of behaviorally relevant stimuli at the single neuron level using a single cell d’ neurometric (d’_sc_),

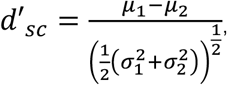

where µ_1_ and µ_2_ were the mean responses of two different stimuli, and σ_1 and_ σ_2_ were the standard deviations of responses to those stimuli.

Population responses (Fig. 3c, Fig. 5d,e), and d’_sc_ profiles (Fig. 3d, Fig. 5d,e) were computed using a moving window (mean +/- SEM) of ΔF values within 10 degree bins in 5 degree increments. Δd’_sc_ profiles were computed using the same procedure with 20 degree bins to increase the number of cells per bin for statistical testing.

To measure the persistence of behavioral bias of each shrew toward novel S-stimuli in the second set of behavioral experiments (Fig. 5f), we least-squares fit each day’s psychometric function (e.g. Fig. 2b, top) with a Gaussian. The shrew’s approximated S+ (the S+ inferred from the shrew’s behavioral performance) was taken as the peak orientation of this Gaussian function (e.g. Fig. 5f, top).

## Acknowledgements

We thank Theo Walker, Val Hoke, and Susan Freling for their assistance with behavioral training and tree shrew care, Dan Wilson and Ben Scholl for expertise with electrophysiological recordings, and Amanda Jacob, Rachel Satterfield, and Nicole Shultz for tissue processing. We also thank members of the Fitzpatrick lab and the MPFI community for support and helpful comments throughout this project. This work was supported by and F32 EY028430 (J.W.S.), R01 EY006821 (D.F.), and Max Planck Florida Institute for Neuroscience.

---

**Extended Data Figure 1:**
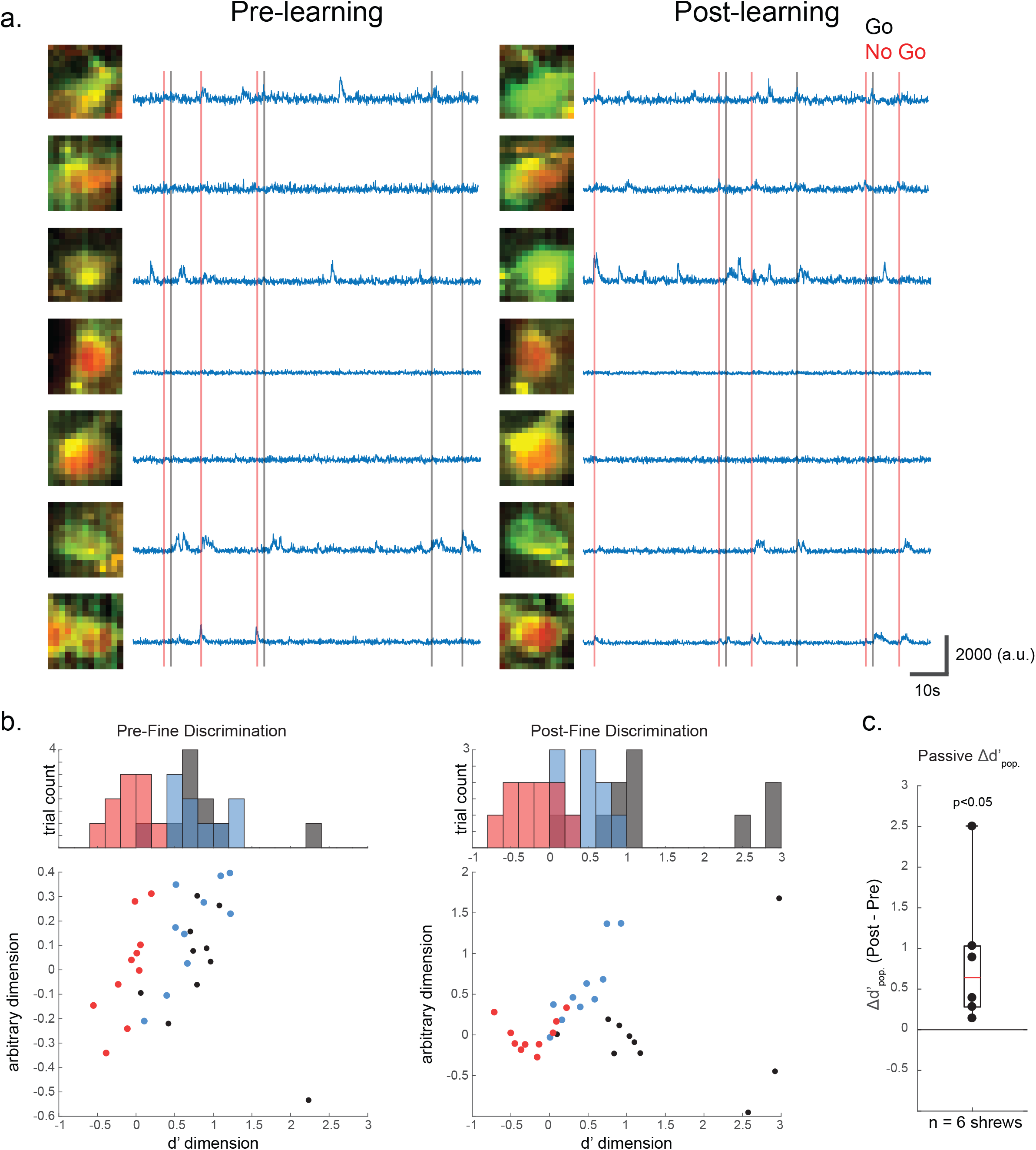
**(a.)** Additional examples of pre- and post-learning raw responses to S+ and S-stimuli in individual V1 neurons. **(b.)** Data from the same animal as in **Figure 2d**, pre-(left) and post-learning (right) clustering of population responses to the S+ (black) 22.5 degree S-(blue) and 45 degree S-(red) shows that in passive stimulus trials, segregation of the population responses in low dimensional space improves with learning. **(c.)** Pre- and post-learning d’_pop_. for 6 imaging subjects demonstrates an increase in population discrimination with learning (p<0.05, Wilcoxon Signed Rank test).

## Notes

### Competing Interest Statement

The authors have declared no competing interest.

